# Mutations of R882 in DNMT3A change flanking sequence preferences and cellular methylation patterns in AML

**DOI:** 10.1101/721472

**Authors:** Max Emperle, Sabrina Adam, Stefan Kunert, Michael Dukatz, Annika Baude, Christoph Plass, Philipp Rathert, Pavel Bashtrykov, Albert Jeltsch

**Author notes:** These authors contributed equally to this work (Max Emperle, Sabrina Adam). Corresponding author: Prof. Dr. Albert Jeltsch, Institute of Biochemistry, Department of Biochemistry, Institute of Biochemistry and Technical Biochemistry, Allmandring 31, D-70569 Stuttgart, Germany.

## Abstract

DNMT3A R882 mutations are frequently observed in AML including the abundant R882H and the rare R882C, R882P and R882S. Using deep enzymology we show here that the DNMT3A-R882H has more than 70-fold altered flanking sequence preferences when compared with wildtype DNMT3A. The R882H flanking sequence preferences mainly differ on the 3’ side of the CpG site, where they resemble DNMT3B, while 5’ flanking sequence preferences of R882H resemble wildtype DNMT3A, indicating that R882H behaves like a DNMT3A/DNMT3B chimera. Activities and flanking sequence preferences of R882C, R882P and R882S were determined as well. Genomic methylation patterns after expression of wildtype DNMT3A and R882H in human cells reflect the flanking sequence preferences. R882H specific hypermethylation in AML patients are correlated with R882H flanking sequence preferences. The hypermethylation events are accompanied by R882H specific misregulation of several genes with strong cancer connection in AML patients, which are potential downstream targets of R882H.

---

DNA cytosine-C5 methylation at CpG sites is a major chromatin regulator essential for development in mammals (Ambrosi et al., 2017; Jeltsch et al., 2018; Schubeler, 2015). DNA methylation is introduced by the family of DNA methyltransferases (DNMTs) in humans comprising DNMT1, DNMT3A, and DNMT3B (Gowher and Jeltsch, 2018; Jeltsch and Jurkowska, 2016; Jurkowska et al., 2011a). It functions in concert with other epigenome modifications, most prominently histone tail modifications and represents one important part of the epigenome network (Allis and Jenuwein, 2016; Soshnev et al., 2018). Aberrant DNA methylation has several connections to diseases including cancer (Baylin and Jones, 2011; Bergman and Cedar, 2013; Feinberg et al., 2016; Norvil et al., 2019). Strikingly, mutations in the DNMT3A enzyme occur in about 25% of all AML patients with a normal karyotype (Hamidi et al., 2015; Yang et al., 2015a). The mutations show a strong enrichment of heterozygous missense mutations (73%) combined with an intact wildtype allele. Among them, R882 mutations are most abundant (about 60%), two third of them R882H, about one third R882C, and 3% R882S and R882P. The high prevalence of the R882H missense mutation, together with the low frequency of nonsense and frameshift mutations in DNMT3A suggests that this DNMT3A mutation has a specific molecular effect, which has been investigated by several groups.

The R882 residue is located at the central interface of the DNMT3A tetramer where it interacts with the DNA backbone (Zhang et al., 2018). Different studies have determined the catalytic activity of the purified R882H mutant protein and mostly observed 50-70% residual activity (Emperle et al., 2018a; Emperle et al., 2018b; Holz-Schietinger et al., 2012; Yan et al., 2011). Different studies reported that the R882H mutation exhibits a dominant negative effect in cells and *in vitro*. Kim et al. (2013) showed that the exogenously expressed murine R878H mutant (corresponding to human R882H) interacts with the wildtype enzyme, but it was less efficient in methylating major satellite repeats in mouse ES cells (Kim et al., 2013). Russler-Germain et al. (2014) described that enzyme preparations obtained after co-expression of wildtype and R882H in mammalian cells showed only 12% of residual activity leading to a model that R882H subunits inactivate the wildtype subunits in mixed complexes in a dominant negative effect (Russler-Germain et al., 2014). However, this model could not be validated biochemically (Emperle et al., 2018a).

It has been well documented that DNMT3A shows pronounced differences in the methylation activity of CpG sites depending on their flanking sequence (Handa and Jeltsch, 2005; Jurkowska et al., 2011b; Lin et al., 2002; Wienholz et al., 2010) and CpG sites have been identified which could not be methylated by DNMT3A at all (Jurkowska et al., 2011b). We observed already in 2006 that the R882A mutation led to a change in the flanking sequence preferences of DNMT3A (Gowher et al., 2006). Recently, we have extended this finding and showed that the flanking sequence preferences of DNMT3A are strongly altered by the R882H mutation leading to specific subsets of CpG sites which are methylated by R882H even better than by wildtype DNMT3A, while another subset of CpG sites shows a drop in activity of R882H that is more pronounced than on average sites (Emperle et al., 2018b). These changes in flanking sequence preferences of R882H may lead to hyper- and hypomethylation of genomic regions and by this play an important role in tumorigenesis by R882H (Emperle et al., 2018b). However, the previous analysis was based on only 56 CpG sites and, therefore, the derived profiles were not sufficiently detailed to correlate them with cellular methylation data.

It was the aim of this work to characterize the flanking sequence differences of DNMT3A and R882H in more detail using a newly developed deep enzymology approach. After confirming and extending the previous findings, we show that other mutations at R882 have similar effects. Moreover, we demonstrate that DNMT3A and R882H transfected in human cells methylate DNA with the same flanking sequence preferences as determined *in vitro*. We demonstrate that specific hypermethylation events occur in AML patients with R882H mutation and they are correlated with the misregulation of several genes with putative connection to AML, which could be part of the tumorigenic downstream signaling cascade emanating from R882H.

## Results

### Generation of flanking sequence preference profiles for DNMT3A and R882H

To determine the sequence preferences of DNA methyltransferases in great depth, we have developed a deep enzymology workflow (Suppl. Fig. 1) (Gao et al., manuscript in preparation). In this approach, a pool of double-stranded DNA substrates is generated which contain one target CpG site flanked by 10 random nucleotides on each side. The substrate pool is enzymatically methylated and after hairpin ligation, bisulfite conversion is carried out which converts cytosine to uracil but leaves 5-methylcytosine intact (Zhang et al., 2009). Afterwards, the bisulfite converted DNA is amplified by two PCRs stepwise adding barcodes and indices (Bashtrykov and Jeltsch, 2018). Different substrates were then mixed and sequenced by NGS. In our previous work, duplicate reactions were carried out with the catalytic domains of DNMT3A and DNMT3B (Suppl. Table 1). Our results demonstrated that DNMT3A- and DNMT3B-mediated methylation reactions show clearly distinct flanking sequence preferences (Gao et al., manuscript in preparation). Here, the specificity of the R882H mutant of DNMT3A catalytic domain was studied with the same approach. Two independent methylation reactions were performed and sequenced at great depth (Suppl. Table 1). To determine the overall influence of each flanking position on the reaction rate, at each site the observed and expected frequencies of each nucleotide in the methylated products were determined. This analysis revealed that the reaction rates of R882H and DNMT3A are both significantly influenced by the CpG-flanking sequence from the −2 to the +3 site (Fig. 1A). Based on this result, we focused on further analyzing the effect of the ±2 bp and the ±3 bp flanking positions on the activity of both enzymes and determined that average methylation for all 256 N2 (NNCGNN) and 4096 N3 (NNNCGNNN) flanks. Correlation analysis of these methylation profiles showed that the repetitions of experiments with the same enzyme always were highly correlated (Fig. 1B). In contrast, the average methylation levels of CpG sites in different flanking context were only weakly correlated between DNMT3A, DNMT3B and R882H confirming the initial observation of altered flanking sequence preference of R882H (Emperle et al., 2018b). To compare the overall methylation activity of R882H with DNMT3A and DNMT3B across all different flanking sites, the DNMT3A results obtained at the lower enzyme concentrations were used and the DNMT3B data were combined. As shown in the heat map in Fig. 1C, both R882H samples are highly similar to each other but dissimilar from DNMT3A and DNMT3B. Therefore, the R882H data were also merged for further analyses and all data normalized to their average methylation.

**Figure 1:**
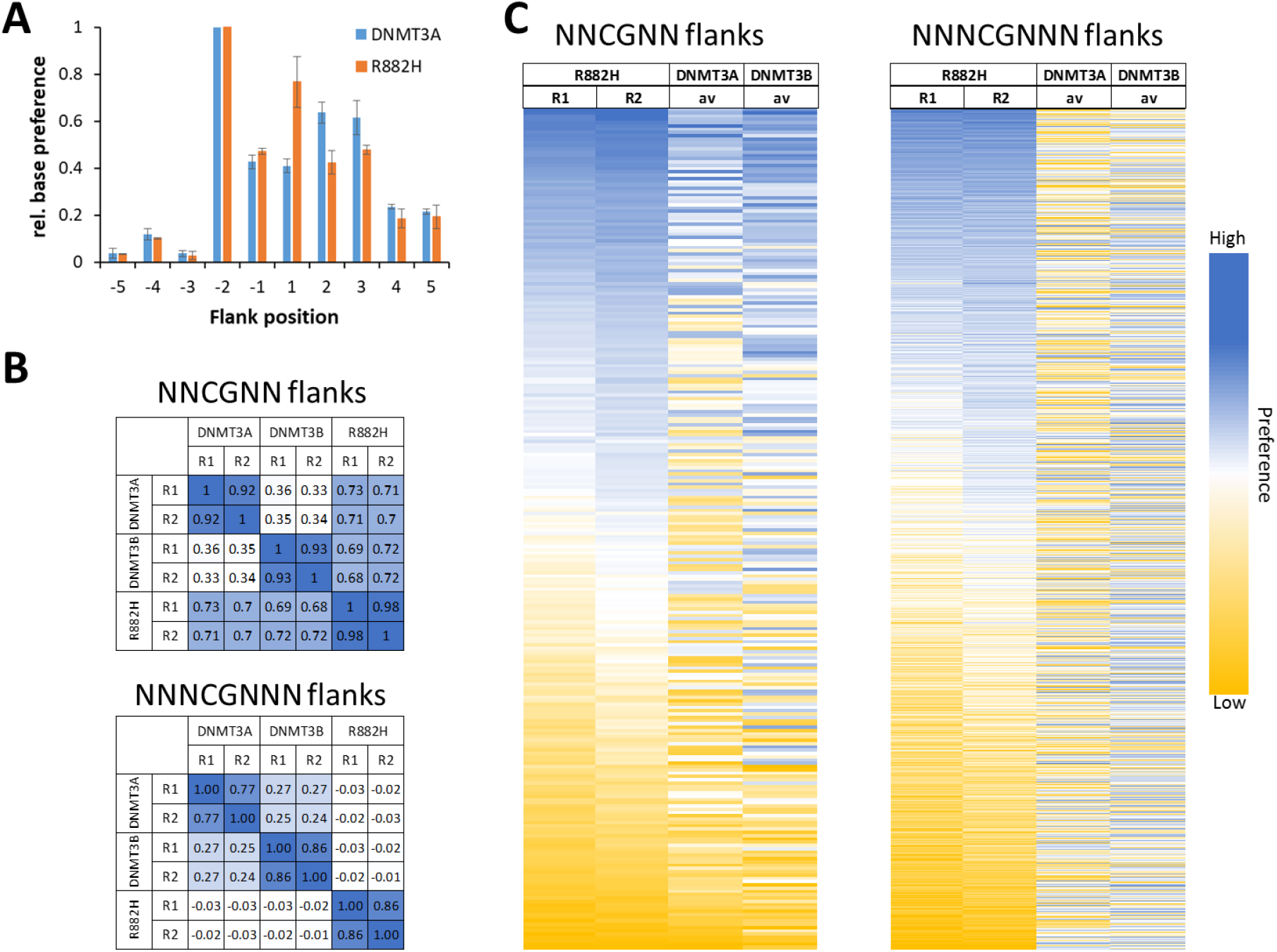
Compilation of the results of the random flank substrate methylation by DNMT3A and R882H. A) Relative base preferences at the −5 to +5 flanking positions of DNMT3A and R882H. The numbers refer to the standard deviations of the observed/expected base composition at each site among the methylated sequence reads, normalized to the highest number for each enzyme. B) Methylation levels were collected and averaged for all two and three base pair flanking sites and the Pearson correlation coefficients (r-values) of the pairwise comparison of the data sets were determined. C) Heatmaps of the methylation profiles of both R882H repeats and the DNMT3A and DNMT3B data. DNMT3A and DNMT3B data were taken from (Gao et al., manuscript in preparation).

Using the averaged and normalized flanking sequence preferences of R882H and DNMT3A, the ratio of R882H/DNMT3A preferences was calculated to express the relative preference of R882H (Fig. 2A). The data revealed a 4.2-fold difference in preferences for NNCGNN flanks. With NNNCGNNN flanks much larger differences in methylation rates were detected. For some of the NNNCGNNN flanks, no methylation was observed with R882H. At these sites the R882H/DNMT3A ratio was set to 0.15 reflecting the limits of detection. Based on this, the difference in preferences was >70-fold for the 3 bp flanks indicating drastic differences in methylation preferences between DNMT3A and R882H. For better visualization Weblogos (http://weblogo.threeplusone.com/) of the 50 most preferred and most disfavored N3 flanks by R882H were prepared (Fig. 2B). These images illustrate a very good agreement of the new flanking sequence preference profiles with the previously published data (Emperle et al., 2018b), although the older data set was based on methylation data of only 56 CpG sites. Specifically, in the previous data set a strong preference of R882H for a G at the +1 and +3 sites was observed together with weaker preference for T at −1 and disfavor for T at the +1 site (Emperle et al., 2018b). All these effects were recapitulated in the detailed analysis based on the deep enzymology data. In the previous publication, two substrates were designed on the basis of the preference profiles available at that time to be preferred and disfavored by R882H to validate the predicted flanking sequence effects (Emperle et al., 2018b). Using these and 7 new substrates designed for different degrees of preference of R882H and DNMT3A (Suppl. Table 2), the rates of methylation of the central CpG site by DNMT3A and R882H were determined to validate the newly derived profiles. After arranging the substrates according to the differences in flanking sequence preferences of R882H and DNMT3A, the relative methylation rates of both enzymes were in a very good agreement with the flanking sequence data (Fig. 2C), indicating that the deep enzymology results are in good agreement with methylation rates determined with these exemplary substrates.

**Figure 2:**
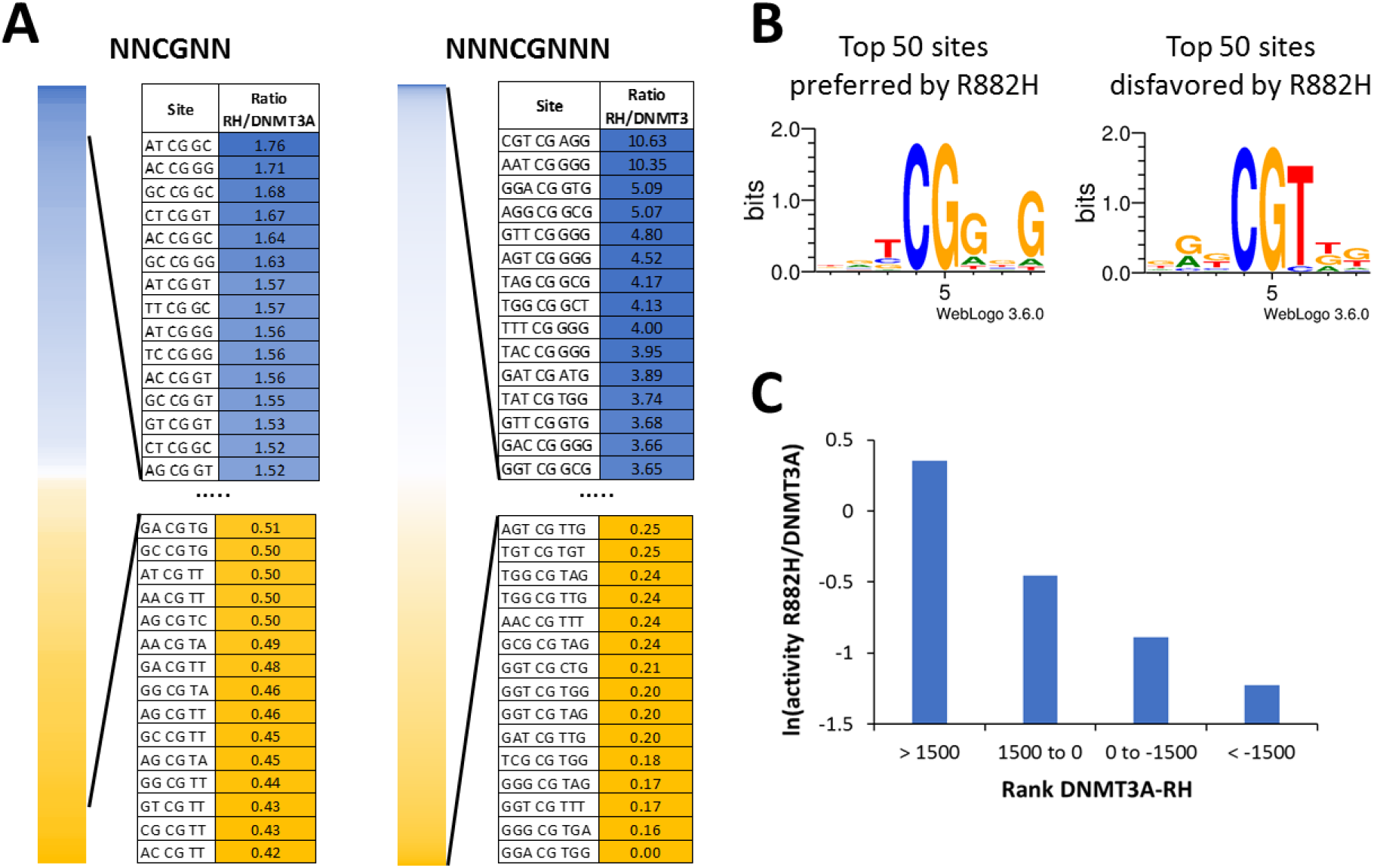
Ratio of methylation rates of R882H and DNMT3A in different flanking context. A) NNCGNN and NNNCGNNN flanking sites ordered by decreasing R882H/DNMT3A preference ratio. B) Sequence logos of the 50 flanking sites most favored or disfavored by R882H. C) Validation of the deep enzymology data by *in vitro* methylation kinetics with 9 defined substrates. Ratios of methylation activities were averaged for different substrates in defined intervals of the difference in the DNMT3A and R882H preference ranks (with a low rank indicating a high preference) (Suppl. Table 3).

### R882H flanking effect on the 5’ and 3’ flank

The structure of DNMT3A/3L in complex with DNA shows that R882 forms a backbone contact to the 3’ flank of the CpG site (Zhang et al., 2018), but it does not contact the 5’ flank (Fig. 3A). Therefore, we were interested to analyze the effect of R882H on the flanking sequence preference separately for the 5’ and 3’ flanks (Fig. 3B). In agreement with the structural data, the 5’ flanking sequence preferences of R882H are highly correlated with those of DNMT3A, while the 3’ preferences differ.

**Figure 3:**
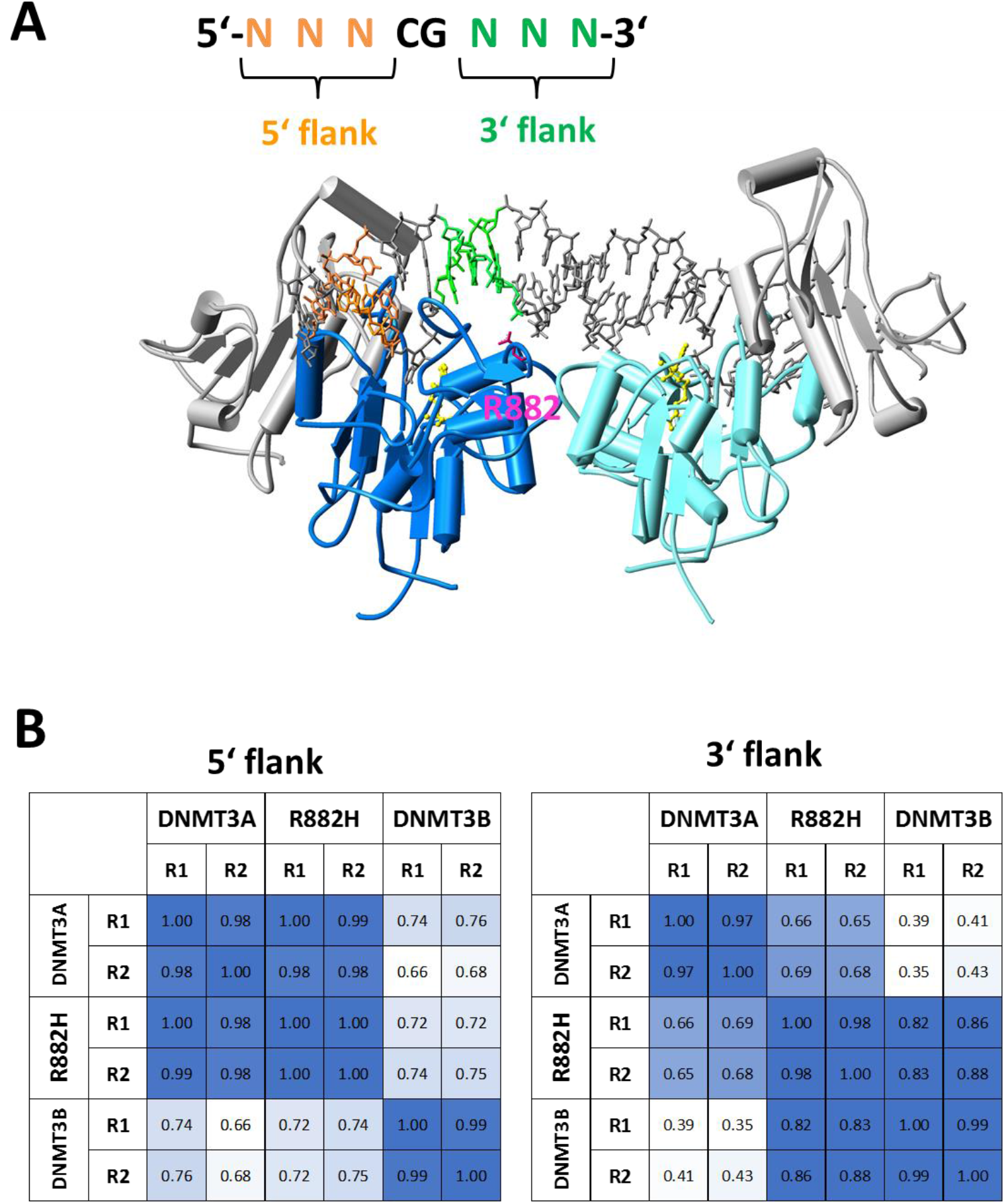
R882H has an effect on the DNA interaction at the 3’ flank. A) Structure of the DNMT3A/3L-DNA complex (Zhang et al., 2018). DNMT3A is shown in dark blue and cyan, DNMT3L in grey. The DNA is shown in grey, with the 5’ flank colored orange and the 3’ flank colored green. R882 is shown for the dark blue DNMT3A subunit in pink. AdoMet is shown in yellow. B) Correlation analysis of flanking sequence preferences of DNMT3A, DNMT3B and R882H for the 5’ and 3’ flanks.

We have shown in our previous paper that the loop containing R882 is a hotspot of amino acid sequence deviation between DNMT3A and DNMT3B, because the DNMT3A residues S881, L883 and R887 are replaced by G822, G824 and K826 in DNMT3B. Moreover, DNMT3B residue R823 (corresponding to R882 in DNMT3A) adopts a different conformation and it does not engage in backbone DNA contact suggesting that this loop region is a major structural cause for the differences in flanking sequence preferences between DNMT3A and DNMT3B (Gao et al., manuscript in preparation). Therefore, it was interesting to compare the DNMT3A and R882H profiles also with DNMT3B (Fig. 3B). The 5’ flank profiles showed high similarity of DNMT3A and R882H, which both differed from DNMT3B. In contrast, the 3’ flank preferences of R882H cluster with DNMT3B much better than with DNMT3A. Strikingly, these data demonstrate that R882H behaves like a chimera of DNMT3A and DNMT3B with respect to flanking sequence preference, it shows DNMT3A preferences at the 5’ flank, but DNMT3B preferences at the 3’ flank. Structurally this result suggests that the loss of the R882 backbone contact in R882H is the main reason for the flanking sequence change.

We also wanted to find out if flanking sequence preferences of DNMT3A, DNMT3B or R882H differ depending on the methylation state of the substrate library pool (unmethylated or hemimethylated). To this end additional deep enzymology reactions were performed with DNMT3A, DNMT3B and R882H in several experimental repeats (Suppl. Table 3). In both settings, methylation of the upper DNA strand was analyzed. We obtained a medium coverage allowing to analyze N2 flanking sequence preferences which revealed that the preferences obtained with unmethylated or hemimethylated substrates were highly similar (Suppl. Fig. 2). Therefore, we were using unmethylated substrates for the further analysis, which are more convenient.

### Investigation of other R882 mutants

As mentioned above, beside R882H also other mutations are occurring at R882 in AML, namely R882C, R882S and R882P. We were interested to find out if these mutations also lead to changes in the flanking sequence preference of DNMT3A. We cloned and purified the mutated proteins in the context of the catalytic domain of DNMT3A. Initially, enzyme activity was analyzed using a 30mer oligonucleotide substrate. Our data showed that R882C and R882P show 30-40% reduction in activity (Fig. 4A), similarly as previously found for R882H with the same substrate (65%) (Emperle et al., 2018a). Interestingly, R882S showed an overall increased activity.

**Figure 4:**
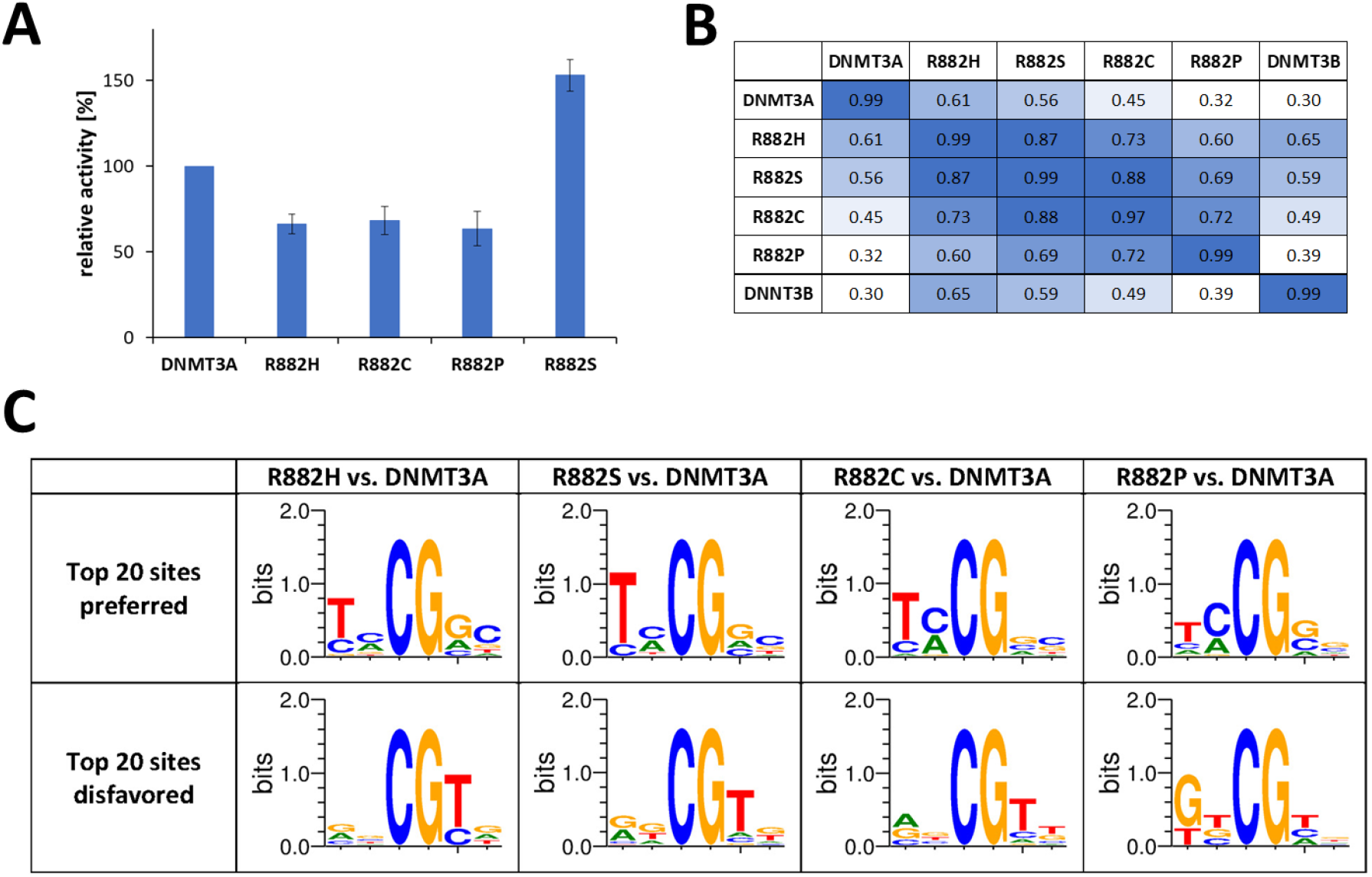
Investigation of other R882 mutants. A) Catalytic activities of R882 mutations determined with a 30mer substrate. The figure shows averages of at least 3 independent experiments. Error bars display the standard deviation. R882H data were taken from Emperle, et al. (2018) (Emperle et al., 2018a). B) Correlation of NNCGNN methylation profiles determined by deep enzymology using the data obtained with unmethylated substrates. For comparison, all data sets were reduced to equal read counts of 38000. C) Weblogos of the 20 most preferred and disfavored flanks by the R882 mutants.

Next, we conducted deep enzymology analyses with the R882C, S and P mutants using unmethylated substrates and obtained a coverage sufficient for the analysis of N2 flanking sequence preferences (Suppl. Table 3). Afterwards, the flanking sequence preferences of all mutants were compared by correlation analysis (Fig. 4B). In addition, Weblogos of the 20 most preferred and most disfavored flanks when compared with wildtype DNMT3A were prepared (Fig. 4C). Strikingly, these analyses showed that the flanking sequence preferences of R882C and R882S are very similar to R882H. This result suggests that the removal of the arginine side chain at position 882and consequently loss of its backbone contact to the DNA is the main cause for the changes in the flanking sequence preferences, rather than specific effects associated with the amino acid introduced instead of arginine at position 882. However, this conclusion is not valid for the R882P mutation, which showed a distinct pattern of flanking sequence preferences that is equally different from DNMT3A, DNMT3B and R882H suggesting some proline specific effects. This result can be rationalized by the strongly reduced regions of allowed ψ and ϕ angles in the Ramachandran plot of proline. Therefore, the introduction of proline in place of an arginine has a high probability to cause larger changes of the loop geometry than all other mutations which would explain unique changes in flanking sequence preferences.

### Analysis of DNA methylation patterns set by DNMT3A R882H in human HCT116 cells

To study the effect of the DNMT3A R882H mutation on DNA methylation in human cells, we used the HCT116 DNMT1 hypomorphic cell line (HCT116 DNMT1 hm), which contains a truncated DNMT1 with reduced activity, but active copies of DNMT3A and DNMT3B (Egger et al., 2006; Rhee et al., 2002). Because of the impaired maintenance DNA methylation activity, these cells have an about 20% reduced level of overall DNA methylation. To facilitate inducible transgene expression, we initially generated a “tet-competent” HCT116 DNMT1 hm cell line via stable integration of the reverse tet transactivator (rtTA3) together with the ecotropic receptor. Subsequently, a retroviral vector was used to generate two cell lines with integrated genes encoding for the full length human DNMT3A-Venus fusion protein or its R882H mutant under the control of a TRE3G promoter (Fig. 5A). A third cell line only expressing the fluorophore served as control. Transgene expression was induced by addition of doxycycline and after 3 days comparable expression levels of DNMT3A and R882H were observed (Suppl. Fig. 3). After 7 days, Venus positive cells were isolated by flow cytometry from all three cell lines. Genomic DNA was isolated and DNA methylation patterns analyzed with Illumina 450k arrays. Heatmap analysis confirmed the similarity of the repeated data sets (Fig. 5B). Filtering of sites showing a significant difference in methylation between wildtype and R882H (p-value < 0.05) indicating that about 80000 sites displayed reduced methylation levels after expression of R882H when compared with cells expressing wildtype DNMT3A (Fig. 5C). Most of them showed up to 20% reduced methylation in agreement with the reduced overall activity of R882H described above. However, about 100 sites showed a stronger (more than 40%) reduction of methylation activity. In addition, more than 9000 sites were identified at which R882H was more active than DNMT3A. Hypermethylation of CpG site in the R882H sample was confirmed at selected sites by amplicon based bisulfite analysis (Fig. 5D). We conclude that altered flanking sequence preferences of R882H are influencing DNA methylation patterns in human cells.

**Figure 5:**
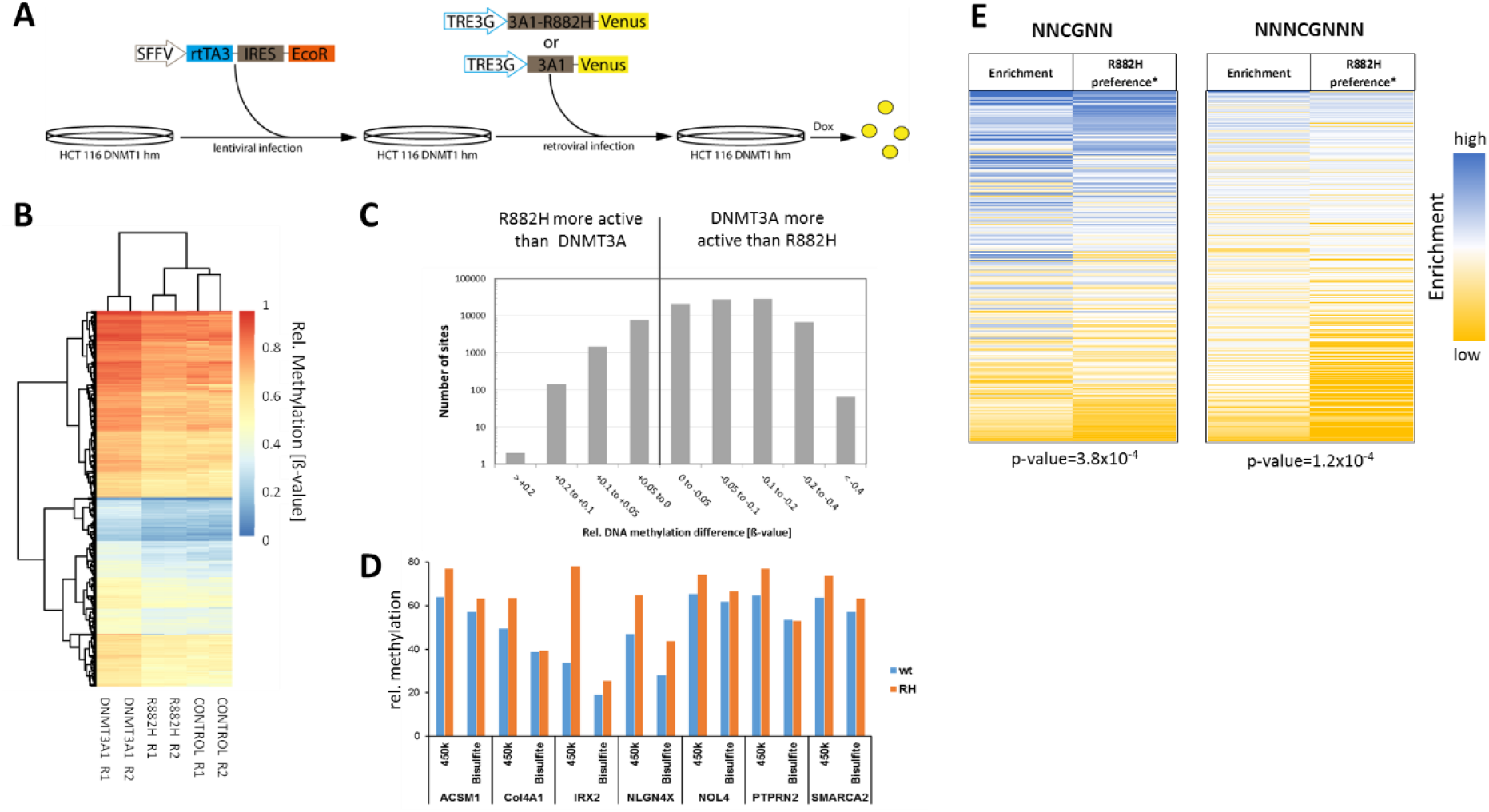
DNA methylation analysis in DNMT1 hypomorphic HCT116 cells. A) Generation of DNMT1 hypomorphic HCT116 cells which express DNMT3A or R882H under a doxycycline (Dox) regulated TRE3G promoter. In the first step, the genomic integration and stable expression of the ecotrophic receptor and the reverse tet-transactivator (rtTA3) was archived using lentiviral infection. In the second step, genomic integration of DNMT3A-Venus or R882H-Venus under the control of a tetracycline-inducible TRE3G promotor was achieved using retroviral infection. Six days after addition of Dox, the cells expressing DNMT3A or R882H were isolated by FACS (Suppl. Fig. 3). B) Clustering of 1000 sites with strongest increase in DNA methylation with the R882H mutant. R1 and R2 denotes the two repeats of the experiment. CONTROL denotes for the untreated cells. C) Overview of all CpG sites showing a significant difference (p<0.05) in the methylation levels of wildtype DNMT3A and R882H. D) Methylation of exemplary CpG sites showing hypermethylation with the R882H mutant on the 450k array was analyzed by bisulfite conversion followed by PCR and amplicon sequencing (Suppl. Table 4). The figure shows the relative methylation of 450k data as ß-values divided by 10 and in % for the bisulfite data. E) Correlation of DNA methylation profiles in cells expressing R882H or wildtype DNMT3A with the flanking sequence preferences of R882H. The enrichment of flanks in 8507 CpG sites that were hypermethylated after expression of R882H in HCT116 cells when compared to cells with expression of wildtype DNTM3A were compared with the R882H/DNMT3A* flanking sequence preference profile. For all flank sizes highly significant correlations were observed. P-values determined by randomization of the data sets are indicated.

When comparing the *in vitro* preference data with cellular data, it needs to be considered that DNMT1 is present in cells, which has a high activity on hemimethylated CpG sites. Therefore, in cells a DNMT3 enzyme needs to methylate just one DNA strand in order to trigger the final appearance of a fully methylated CpG site. To account for this, we have defined symmetrical R882H/DNMT3A preference profiles by averaging the preferences of pairs of corresponding complementary flanks (called R882H/DNMT3A* preference) (Suppl. Fig. 4). To correlate the R882H specific hypermethylation profiles in HCT116 DNMT1 hm cells with the R882H/DNMT3A* preference, all hypermethylated sites with P<0.05 and at least 0.5-fold higher methylation in the R882H data set were selected (8507 CpGs). Among this set, the enrichment of two and three bp flanks was determined when compared with the distribution of flanking sites among the entire 450k data set. The overrepresentation of flanks in the hypermethylated data set was compared with the R882H/DNMT3A* preference (Fig. 5E) revealing clear correlations between the enrichment of flanks in the hypermethylated data set with the R882H/DNMT3A* flanking sequence preferences for all flank lengths. To determine the statistical significance of these correlations, the correlation analyses were repeated with randomly shuffled enrichment data. Based on the distribution of the r-values in these random correlations, p-values were determined, indicating that all correlations are highly significant with p-values of 3.8×10^−4^ for N2 flanks and 1.2×10^−4^ for N3 flanks, respectively. These results clearly indicate that DNA methylation patterns generated in human HCT116 DNMT1 hm cells by wildtype DNMT3A and the R882H mutant reflect the flanking sequence preferences of both enzymes, indicating that the flanking sequence preferences influence the DNA methylation activity of DNMT3A in human cells.

### Correlation of major satellite methylation with the R882H/DNMT3A* flanking sequence preferences

As described above Kim et al. (2013) demonstrated a dominant negative effect of R882H expression on DNA methylation of major satellite repeat sequences in murine ES cells (Kim et al., 2013). They conducted DNA methylation analysis by Southern blot hybridization after digestion of the DNA with the methylation specific restriction enzyme MaeII (ACGT) probing 3 CpG sites and by bisulfite analysis of 6 CpG sites. We investigated the preference of R882H for these sites and strikingly observed that the majority of the sites on the major satellite were strongly disfavored by R882H (Suppl. Fig. 5). Hence our data are in good agreement with the experimental observations reported in this paper.

### Correlation of AML patient DNA methylation profiles with the R882H* flanking sequence preferences

Next we aimed to compare global DNA methylation patterns from AML patients carrying the R882H mutation with AML patients not containing it. Glass et al. (2017) reported enhanced reduced representation bisulfite sequencing (ERRBS) DNA methylation profiles of 119 AML samples, 27 of them with R882H mutation (Glass et al., 2017). The data include CpGs covered between 10-400x, methylation changes were called requiring at least 3 samples to support the difference. In total 184589 differentially methylated CpGs with a q-value ≤0.01 and ≥25% methylation change were identified for AML with R882H mutations as compared to AML with wildtype DNMT3A, 56870 of them were hypermethylated in R882H AML and 127719 were hypomethylated (Glass et al., 2017). The methylation levels of the sites specifically hypermethylated in R882H containing AMLs were binned into different categories depending on the degree of hypermethylation (Suppl. Fig. 6) and for each group the distribution of NNCGNN flanks was determined. Next, the frequency of flanks within certain preference ranges of R882H were determined in all groups (Fig. 6A). Strikingly, the Top-500 group containing the 500 sites most strongly hypermethylated in R882H containing AML across the entire data set of 119 AML cases showed a very strong enrichment of flanks which are highly preferred by R882H, while flanks disfavored by R882H were depleted. A similar but less pronounced effect was observed in the Top-1000 group. The strong enrichment of R882H preferred flanks in the Top-500 and Top-1000 groups is also clearly visible in the corresponding heatmaps (Fig. 6B). To determine the statistical significance of these correlations, the correlation analyses were repeated with randomly shuffled R882H* data. Based on the distribution of the r-values in these random correlations, p-values were determined, indicating that the correlations observed with the Top-500 (p-value 3.5×10^−5^) and Top-1000 (p-value 4.4×10^−4^) groups are highly significant (Fig. 6C). This finding clearly indicates that the altered flanking sequence preferences of the R882H mutant shape the global DNA methylation profile in AML patients. It is very striking that these effects were detectable in a global data set obtained by averaging 119 AML patient data (27 of them with R882H mutation and 92 without) despite the high background of DNA methylation changes related to AML progression (Spencer et al., 2017). These data suggest that corresponding alterations in the DNA methylation might cause disease relevant expression changes of genes with oncogenic or tumor suppressive roles.

**Figure 6:**
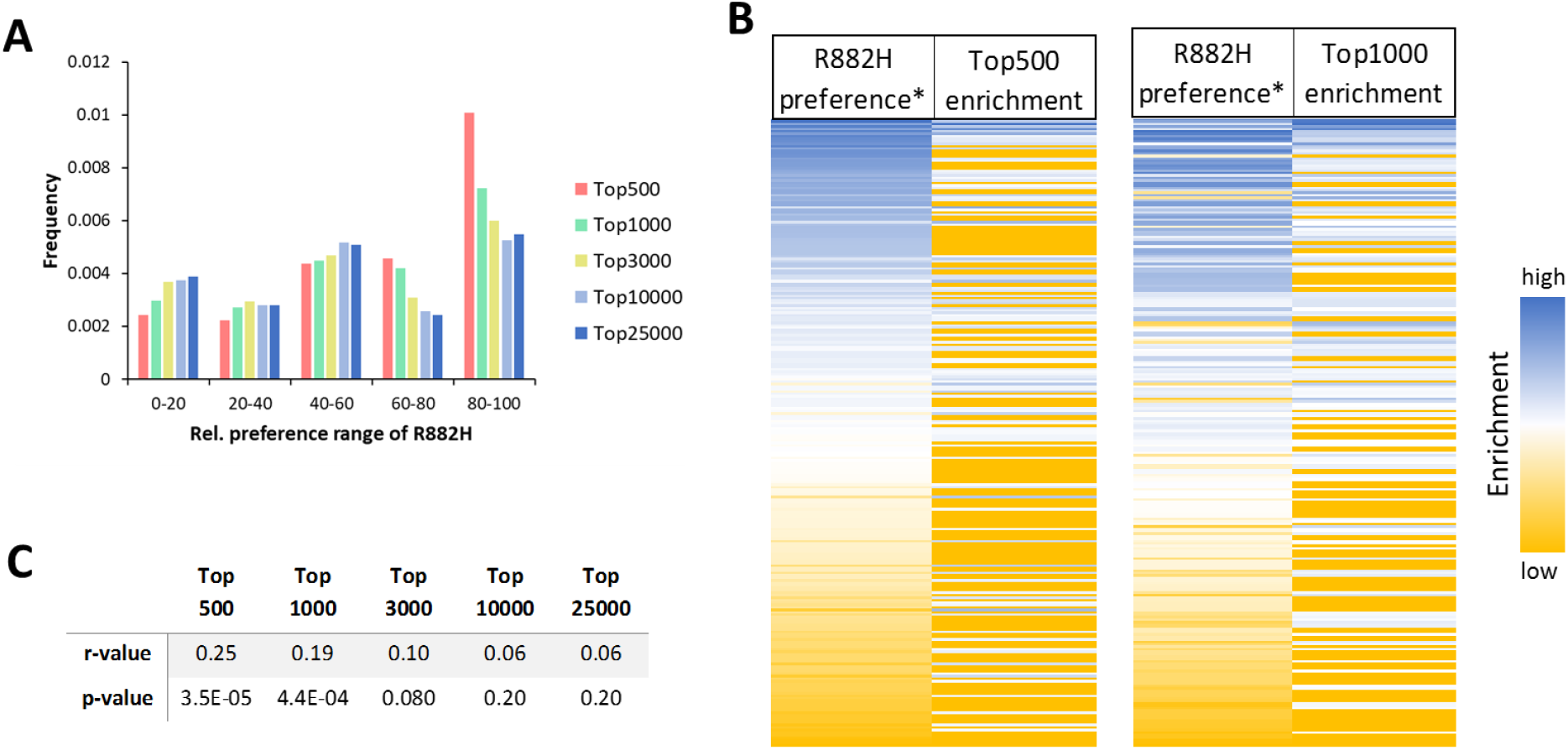
Correlation of R882H flanking sequence preference with the flanking sequence context of CpG sites specifically hypermethylated in R882H containing AML (Glass et al., 2017). A) Enrichment or depletion of R882H*preferred flank in the different hypermethylated groups. Flanks strongly preferred by R882H have a relative preference of 100 disfavored flanks have a preference of 0. B) Correlation of the enrichment of flanks in the Top500 and Top1000 group with the R882H* preference (RH*). C) Compilation of r-values and corresponding p-values of the correlation of the R882H flanking sequence preferences with the enrichment of flanking in the different groups of hypermethylated sites.

### Identification of potential AML target genes regulated by R882H

Next we aimed to identify if DNA hypermethylation in the R882H data set was correlated with R882H specific expression changes reported by Glass et al. (2017). For this we focused on the hypermethylated CpG sites which were assigned to gene promoters, gene bodies or close to genes with a distance of 2-50 kb. The rationale behind this approach was that hypermethylation of gene promoters often leads to repression of genes, but some transcription factors have also been identified to preferentially bind methylated CpG sites (Kribelbauer et al., 2017; Yin et al., 2017). Hypermethylation of gene bodies often increases gene expression (Su et al., 2018; Yang et al., 2014). In addition, methylation of enhancers in gene bodies and close to genes can influence expression in both directions, depending on the effect of the regulated transcription factors (stimulatory or repressive) and their methylation specificity. In total 136 genes were identified in which R882H specific DNA hypermethylation was co-occurring with an R882H specific expression change. These initial targets were further analyzed for a potential role in cancer and AML in particular, finally leading to a list of 17 top-candidates, several of them associated with more than one hypermethylated CpG site (Table 1). 12 of the genes contain at least one hypermethylated CpG site in flanking context that is among the 10% most preferred flanks of the R882H* profile, the remaining 5 genes contained a CpG in the 40% preferred context. Many of the downregulated genes were known to be tumor suppressor genes and many of the upregulated ones oncogenes. The strong correlation of PRDM16 over-expression with DNMT3A mutations has already been reported (Yamato et al., 2017) and PRDM16 is a known oncogene and its expression is correlated with poor clinical outcome in AML (Shiba et al., 2016). Overexpression of WT1 is generally observed in AML and it has been associated with poor clinical outcome in many studies (Rampal and Figueroa, 2016). Additional literature evidence suggests roles of TP53INP1 and PER3 in AML, because down-regulation of any of them is a negative prognostic marker in AML (Torrebadell et al., 2018; Yang et al., 2015b). We conclude that these genes represent strong candidates for misregulated targets downstream of R882H which could have a direct influence on AML onset and progression.

**Table 1:**
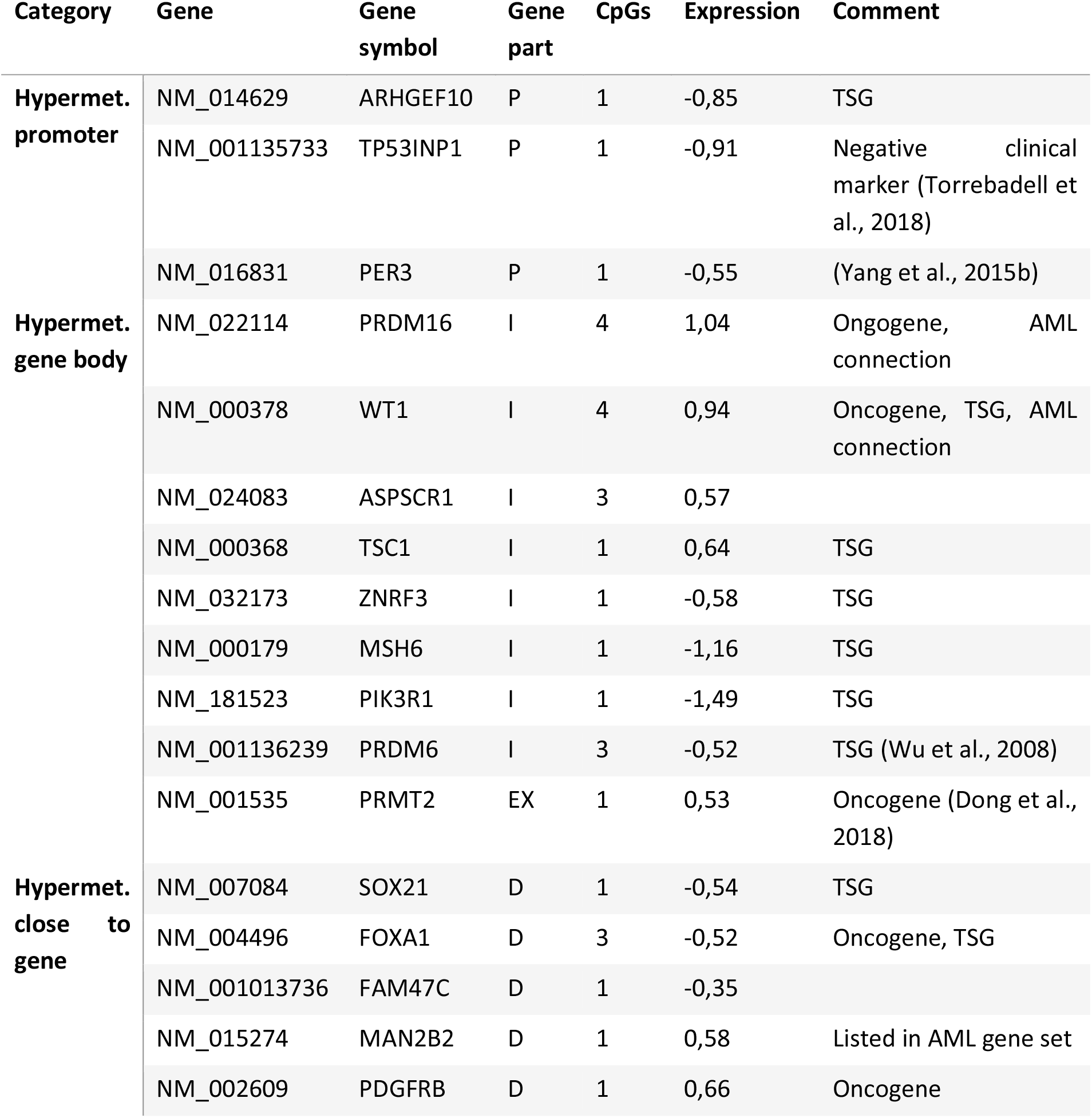
List of potential R882H AML target genes. Gene part refers to the region of hypermethylation (P – promoter, I – Intron, EX – Exon, D – Distance) as defined in (Glass et al., 2017). CpGs: number of hypermethylated CpGs, Expression: ln2 ratio of expression level in R882H and non-R882H AML. Comment: Oncogene and TSG (Tumor suppressor gene) as listed in COSMIC Cancer gene census (Sondka et al., 2018) or the indicated references. AML connection is based on Harmonizome AML gene set (Rouillard et al., 2016).

## Discussion

Cancer is caused by genetic mutations and epigenetic changes (epimutations) which in combination lead to the characteristic physiological and morphological features of tumor cells, most importantly unregulated cell division. These effects are connected in several instances, for example mutations or epigenetic silencing of DNA repair genes leads to a mutator phenotype stimulating the generation of genetic changes in tumor cells and, conversely, the mutation of epigenetic factors can perturb the epigenetic signaling system leading to the increased occurrence of epimutations (Feinberg et al., 2016; Plass et al., 2013). One example of the latter case are mutations in the DNA methyltransferase DNMT3A which are observed in about 25% of all AML patients with normal karyotype. About 60% of all missense mutations in DNMT3A occur at R882 and in about 2/3 of them are R882H exchanges. The high frequency of this specific mutation has stimulated several studies to investigate the pathogenic mechanism of R882 mutations, in particular R882H, but its role has not been finally settled. Previous studies have provided evidence for a dominant negative effect of R882H (Russler-Germain et al., 2014), but this result has remained controversial (Emperle et al., 2018a). We have recently demonstrated that the R882H mutation causes a strong shift in the flanking sequence preference of DNMT3A affecting 3 base pairs of DNA sequence surrounding the CpG target site (Emperle et al., 2018b). This finding was in agreement with structural data showing that R882 is located at the DNA binding site of the enzyme where it forms a backbone phosphate contact to the flanking base pair at the +3 position (Zhang et al., 2018). However, the flanking sequence preference data in the previous paper were based on a limited set of kinetic data referring to only 56 different CpG sites. Therefore, these data were insufficient for the correlation of R882H specific changes of flanking sequence preferences with cellular methylation pattern which would have to consider target sites embedded into 4096 different 3 base flanking context (NNNCGNNN).

In this work we have exploited a recently developed deep enzymology approach to analyze flanking sequence preferences of R882H and other R882 mutations in great detail. In this technique, a pool of DNA substrates is generated in which the CpG target site is embedded in a 10 base pair random sequence context on either side. The mixture of substrates is methylated and methylation levels determined by bisulfite conversion coupled to NGS. Depending on the read depth, average methylation levels of all N2 or N3 flanking sequences can be determined, allowing to derive detailed flanking sequence preference profiles. We previously conducted deep enzymology flanking sequence analysis for DNMT3A and DNMT3B (Gao et al., manuscript in preparation) and now added detailed data on R882H. The new results clearly confirm the previous observation of altered flanking sequence preferences of R882H (Emperle et al., 2018b) now showing up to 70-fold differences in the methylation rates of CpG sites in different N3 flanking environments. Strikingly, a comparison of the flanking sequence preferences of R882H with DNMT3A and DNMT3B revealed that R882H behaves like a chimera of DNMT3A and DNMT3B. With respect to the 5’ flank, the flanking sequence preferences of R882H and DNMT3A were very similar and distinct from DNTM3B. At the 3’ flank, however, the profiles of R882H and DNMT3A differed and R882H was more similar to DNMT3B. This result is in agreement with our previous finding, that the loop containing R882 is one key determinant for the divergence of the flanking sequence preferences of DNMT3A and DNMT3B (Gao et al., manuscript in preparation). We also prepared the R882C, S and P mutations and investigated their catalytic activity showing that the activities of R882C and R882P are similar to R882H, but R882S is even more active than wildtype DNMT3A, arguing against a loss-of-function mechanism for this mutation. We determined the flanking sequence preferences of all three mutants showing that the profiles of R882S and R882C are very similar to R882H, suggesting a common pathomechanism. This result also indicates, that the shift in the flanking sequence preferences most likely is caused by the loss of the arginine at positon 882 and its contact to the DNA, and it is not due to the presence of the new amino acid at this site. In case of R882P, the flanking sequence preferences were equally weakly correlated with DNMT3A, R882H and DNMT3B, suggesting some specific effects of the introduced proline.

Next we aimed to compare the *in vitro* flanking sequence preferences of R882H with cellular methylation data. To this end, symmetrical R882H/Dnmt3A* preference profiles were calculated and compared with methylation data obtained after expression of DNMT3A and R882H in HCT116 cells containing a hypomorphic DNMT1. A clear and highly significant correlation was observed demonstrating that flanking sequence preferences also guide the activity of DNMT3A and R882H in human cells. We also noticed a clear correlation of the detailed flanking sequence preferences of R882H reported here with published DNA methylation data in mice showing that the expression of R882H in mouse ES cells reduced the methylation of major satellite repeats (Kim et al., 2013). Strikingly, the flanking sequence preferences of R882H readily explain this finding, because the CpG sites present in the mouse major satellite repeats are strongly disfavored by R882H. We next aimed to analyze patient DNA methylation data to determine if an effect of the altered flanking sequence preferences of R882H is detectable in them as well. To this end, we used a comprehensive analysis of DNA methylation profiles of 119 AML patients, 27 of them with R882H (Glass et al., 2017). Using the CpG sites consistently hypermethylated in R882H containing tumors, we find that the sites with strongest hypermethylation show a striking and statistically highly significant correlation with R882H flanking sequence preferences, indicating that even in this highly aggregated analysis, the specific flanking sequence preferences of the R882H mutation are still detectable. This result strongly suggests that the flanking sequence preferences of DNMT3A and R882H are an important determinant of the global genome wide DNA methylation patterns in AML cells. To investigate potential downstream effect of the R882H induced hypermethylation, we combined the DNA methylation data with gene expression data (Glass et al., 2017) and functional gene information and compiled a list of 17 genes with strong tumor connection that are connected to R882H specific hypermethylated CpG sites and that show R882H specific changes in gene expression. These genes are strong downstream target candidates potentially misregulated by R882H and then promoting tumor formation.

## Conclusions

In this work we describe the biochemical mechanism of a gain-of-function effect caused by the R882H mutation in DNMT3A. By changing the flanking sequence preferences of DNMT3A the mutation leads to altered DNA methylation patterns in human cells and patient samples, including the hypermethylation of about one third of the differentially methylated CpG sites. While our data are not excluding specific effects of hypomethylated CpG sites, an important role of the hypermethylated ones is in agreement with the genetic finding of the massive enrichment of heterozygous R882H point mutations in patients and the co-occurrence of them with IDH and TET2 mutations, both leading to reduced DNA demethylation. By data mining of patient data we identify a group of with strong cancer connection, which are specifically hypermethylated and misregulated in R882H AML. These genes are strong downstream target candidates of R882H and they represent potential targets for further treatment of R882H containing AML.

## Methods

### Biochemical methods

For biochemical studies, the C-terminal domain of human DNMT3A (amino acids 612–912 of Q9Y6K1) and its R882H mutant and DNMT3B (amino acids 558-859 of O88509) were used. All were cloned into pET28+ (Novagen) containing an N-terminal His6-tag. Mutagenesis was performed using the megaprimer site-directed mutagenesis method (Jeltsch and Lanio, 2002) and confirmed by restriction marker analysis and DNA sequencing. Protein expression and purification was performed as described (Emperle et al., 2014). Methylation activity was determined using radioactively labelled AdoMet (Perkin Elmer) and a biotinylated double stranded 30mer oligonucleotide as described (GAG AAG CTG GGA CTT CCG GGA GGA GAG TGC) (Emperle et al., 2018a). Specific methylation of one CpG site in the upper DNA strand was determined using hemimethylated (with methylation in the lower strand of the CpG site) oligonucleotide substrates. In addition, the methylation of the same substrate in fully methylated state was determined and the rates subtracted from the methylation rates of the hemimethylated substrates as described (Emperle et al., 2018b; Jurkowska et al., 2011b). This approach allows to detect the specific methylation of the target site ignoring additional methylation events in the remaining part of the substrate. This special DNMT assay is necessary for DNMT3 enzymes if methylation of individual sites should be measured, because of their high non-CpG methylation activity (Gowher and Jeltsch, 2001).

### Deep enzymology experiments

Deep enzymology experiments were conducted as described (Gao et al., manuscript in preparation). In short, libraries of DNA substrates containing unmethylated and hemimethylated CpG sites embedded in a 10 nucleotide random context were prepared by primer extension using single stranded templates obtained from IDT. The pool of substrates was methylated at 37 °C in the presence of 0.8 mM S-adenosyl-L-methionine (Sigma) in reaction buffer (20 mM HEPES pH 7.5, 1 mM EDTA, 50 mM KCl, 0.05 mg/mL bovine serum albumin) using enzyme concentrations and incubation times as indicated in the text. Stopping of the methylation was followed by hairpin ligation and bisulfite conversion using EZ DNA Methylation-Lightning kit (ZYMO RESEARCH). Libraries for Illumina Next Generation Sequencing (NGS) were produced with the two-step PCR approach pooled in appropriate ratios and analyzed by Illumina NGS.

### Bioinformatics

Bioinformatics analysis of NGS data was conducted with the tools available on the Usegalaxy.eu server (Afgan et al., 2018) and with home written programs. First, fastq files were analyzed by FastQC, 3’ ends of the reads with a quality lower than 20 were trimmed and reads containing both full-length sense and antisense strands were selected. Next, using the information of both strands of the bisulfite-converted substrate the original DNA sequence and methylation state of both CpG sites was reconstituted and average methylation levels of each NNCGNN and NNNCGNNN site were determined. Pearson correlation factors were calculated with Excel using the correl function. P-values were determined using the distribution of r-values from >200 correlation analyses with one data set shuffled. ERRBS DNA methylation data for 119 AML patients with annotated R882H mutational status were obtained from GSE86952 (Glass et al., 2017). Average NCGN, NNCGNN and NNNCGNNN methylation levels in BED file obtained from 450k analyses and ERRBS data were determined using home written programs.

### Generation of HCT116 cells expressing DNMT3AC and R882H

HTC116 DNMT1 hypomorphic cells (HCT116 DNMT1 hm) (kindly provided by Prof. Bert Vogelstein, HHMI, USA), were cultured in McCoy 5A Medium (Sigma) supplemented with 10% FCS, 2 mM L-glutamine, 100 U/ml penicillin and 100 μg/ml streptomycin at 37°C in 5% CO_2_. HTC116 DNMT1 hm cells were modified to express the ecotropic receptor and rtTA3 using retroviral transduction of pWPXLd-RIEP (pWPXLd-rtTA3-IRES-EcoR-PGK-Puro) followed by drug selection (0.8 ug/ml puromycin for 1 week, respectively) similarly as described (Rathert et al., 2015). The resulting cell line was subsequently transduced with ecotropically packaged retroviruses containing the *dnmt3a* (or R882H mutant) gene fused to Venus under control of a TRE3G promoter. Retroviral gene transfer was performed as previously described (Rathert et al., 2015). In brief, for each calcium phosphate transfection, 10‐20 μg plasmid DNA and 5 μg helper plasmid (pCMV-Gag-Pol, Cell Biolabs) were used. Retroviral packaging was performed using PlatinumE cell line (Cell Biolabs). Transduction efficiencies of retroviral constructs (TRE3G-DNMT3A1-Venus-PGK-NEO, TRE3G-DNMT3A1(R882H)-Venus-PGK-NEO or TRE3G-Venus-PGK-NEO) (Liu et al., 2014) were measured 48h post induction with 1 μg/ml doxycycline (DOX) by flow cytometry. Single‐copy conditions were achieved by infecting less than 20% of the initial population, guaranteeing <2% cells with multiple integrations. Transduced cell populations were selected 48 h post infection using 500 μg/ml G418 (Gibco Life technologies). After 7 days of induction about 1 million DNMT3A or R882H expressing cells (as judged by being Venus^+^) were sorted for each replicate by FACS.

### Infinium 450k DNA methylation analysis

To determine genome wide DNA methylation of HCT116 DNMT1 hm cells expressing DNMT3A wildtype or R882H, the Infinium 450k array was used according to the instructions of the DKFZ Genomics and Proteomics Core Facility. Two repeats of each data set were prepared and used for statistical analysis. From the raw intensity data (in total 485,577 sites), 35654 CpG sites were removed during quality control, the remaining were background-subtracted with the methylumi (R package version 2.18.2) and normalized by the BMIQ method. Additional 1416 CpG sites were removed from analysis after normalization. For further analysis, the methylation levels of R882H and wildtype were compared and 92823 sites with significant difference in methylation (p-value <0.05) were extracted. In the next step, R882H hypermethylated sites were ranked by comparing the relative and absolute methylation difference, and the methylation data of the 1000 most significant sites were clustered (Euclidean distance, complete linkage) and visualized using pheatmap (Kolde R, R package version 1.0.8).

### Amplicon-targeted bisulfite sequencing

To determine the methylation levels of selected CpG sites, the DNA samples used for 450k analysis were digested overnight at 37°C using BamHI and EcoRI and then bisulfite converted using EZ DNA Methylation-Lightning Kit (Zymo Research) following the manufacturer’s instructions. The bisulfite treated DNA was then used for PCR amplification using the amplicon and primers with treatment specific barcodes and HotStarTaq DNA Polymerase (Qiagen). PCR products were resolved on an 1% agarose gel, followed by gel extraction and clean-up using the NucleoSpin Gel and PCR Clean-up (Macherey-Nagel). The products were mixed at an equimolar ratio and sent for paired-end sequening on Illumina HiSeq2000 to Novogene Bioinformatics Technology Co., Ltd., Beijing, China (www.novogene.cn). The high-throughput sequencing results were demultiplexed and analyzed using the CLC Genomic Workbench 10.0.1 (CLC Bios, MA) following the manufacturer’s standard data import protocol and the bisulfite sequencing plugin and the methylation levels normalized to the total read number where extracted.

## Acknowledgements

We gratefully acknowledge sharing of primary expression data published in Glass et al. (2017) by Drs. Maria Figueroa and Jacob Glass. This work has been supported by the Deutsche Forschungsgemeinschaft grants JE252/20 and JE252/36 and the Sander Foundation.

