## Supplemental Tables and Figures for "Mutations of R882 in DNMT3A change flanking sequence preferences and cellular methylation patterns in AML"

### Supplemental figures and tables

##### Suppl. Tables

Suppl. Table 1: NGS results of the DNMT3A, DNMT3B and R882H samples using hemimethylated substrates.

Suppl. Table 2: List of the methylation substrates used for the validation of the deep enzymology data.

Suppl. Table 3: NGS results of the DNMT3A, DNMT3B and different R882 mutants with medium coverage using unmethylated substrates.

Suppl. Table 4: List of the bisulfite primers used for validation of the 450k array data.

##### Suppl. Figures

Suppl. Fig. 1: Deep enzymology workflow allowing the methylation of DNA substrates in randomized $N_{10}CGN_{10}$ libraries.

Suppl. Fig. 2: Correlation analysis of NNCGNN deep enzymology data obtained for DNMT3A, DNMT3B and R882H with hemimethylated (HM) and unmethylated (UM) substrates.

Suppl. Fig. 3: FCS profile of DNMT1 hypomorphic HCT116 cells with stable insertion of the DNMT3A (3A1) gene or its R882H mutant (3A1-R882H).

Supplemental Fig. 4: Definition of R882H/DNMT3A* preference profile as the average of the preferences of reverse complementary sites.

Supplemental Fig. 5: Ranking of the flanks of CpG sites analyzed by Kim et al. (2013) to study major satellite methylation in murine ES cells in the R882H* preference (Kim et al., 2013).

Suppl. Fig. 6: Binning of the 56580 sites that were specifically hypermethylated in R882H containing AML in the data set of Glass et al. (2017) into different tiers depending on their degree of hypermethylation (Glass et al., 2017).

**Suppl. Tables**

Suppl. Table 1: NGS results of the DNMT3A, DNMT3B and R882H samples using hemimethylated substrates. Incubation times were 60 min in each case. The results of DNMT3A and DNMT3B were already presented in (Gao et al., manuscript in preparation).

| Sample | $c_{enzyme}$ [µM] | No. of reads |
|---|---|---|
| DNMT3A R1 | 0.5 | 653331 |
| DNMT3A R2 | 1 | 131110 |
| DNMT3B R1 | 2 | 93991 |
| DNMT3B R2 | 4 | 951779 |
| R882H R1 | 2 | 169532 |
| R882H R2 | 4 | 1260190 |

Suppl. Table 2: List of the methylation substrates used for the validation of the deep enzymology data. All flanking sites were embedded into the 30mer substrate used before (Emperle et al., 2018a). Data for substrate 1 and 4 were taken from (Emperle et al., 2018b).

| Name | Sequence | Rank DNMT3A | Rank R882H | Rank DNMT3A-R882H | Rel. activity R882H/DNMT3A |
|---|---|---|---|---|---|
| Substrate 1 | AAC CG GTG | 3113 | 439 | 2674 | 1.42 |
| Substrate 2 | TTC CG GGA | 909 | 624 | 285 | 0.64 |
| Substrate 3 | TAT CG TGG | 3947 | 4050 | -103 | 1.05 |
| Substrate 4 | ACC CG GGG | 2837 | 3250 | -413 | 0.46 |
| Substrate 5 | GGT CG TAA | 2420 | 3171 | -751 | 0.14 |
| Substrate 6 | AGG CG CCC | 1220 | 1935 | -715 | 0.07 |
| Substrate 7 | TTA CG CCC | 49 | 145 | -96 | 0.30 |
| Substrate 8 | GTA CG CCA | 125 | 1506 | -1391 | 0.43 |
| Substrate 9 | GTA CG TCA | 57 | 2591 | -2534 | 0.29 |

Suppl. Table 3: NGS results of the DNMT3A, DNMT3B and different R882 mutants with medium coverage using unmethylated substrates. For further analysis the individual repeats of the DNMT3A, DNMT3B and R882H samples were combined.

| Sample | $c_{enzyme}$ [µM] | Incubation time [min] | No. of reads |
|---|---|---|---|
| 3A-UM-R1 | 0.5 µM | 60 min | 56344 |
| 3A-UM-R2 | 1 µM | 60 min | 44676 |
| 3B-UM-R1 | 1 µM | 30 min | 53367 |
| 3B-UM-R2 | 1 µM | 60 min | 55188 |
| 3B-UM-R3 | 2 µM | 30 min | 56035 |
| 3B-UM-R4 | 2 µM | 60 min | 57577 |
| R882H-UM-R1 | 0.5 µM | 60 min | 35709 |
| R882H-UM-R2 | 2 µM | 60 min | 34807 |
| R882C | 0.5 µM | 60 min | 47648 |
| R882P | 0.5 µM | 60 min | 41655 |
| R882S | 0.5 µM | 60 min | 48626 |

Suppl. Table 4: List of the bisulfite primers used for validation of the 450k array data.

| Name | Primer sequence |
|---|---|
| ACSM1_bis_f | TAGGTGGTGATTTGAGAATTTTGTG |
| ACSM1_bis_r1 | AGTACTTCCATACCTTTCAACCTAATAAAAACA |
| ACSM1_bis_r2 | GATCGTTCCATACCTTTCAACCTAATAAAAACA |
| COL4A1_bis_f | GTGAGATGATGGTTAATGGTTTGTTT |
| COL4A1_bis_r1 | AGTACTCTTTTTCTCCTTCAACAAATAAAAATC |
| COL4A1_bis_r2 | GATCGTCTTTTTCTCCTTCAACAAATAAAAATC |
| IRX2_bis_f | TTGGGTTTGGGTTTAGGGTAGTTA |
| IRX2_bis_r1 | AGTACACAAAAAATCCACTTTACTTTTAACCTC |
| IRX2_bis_r2 | GATCGACAAAAAATCCACTTTACTTTTAACCTC |
| NLGN4X_bis_f | TGGTTTTTGTATTTTTTTGGATGAG |
| NLGN4X_bis_r1 | AGTACAAAACTTTCCATCCTTTACTACAAATCAA |
| NLGN4X_bis_r2 | GATCGAAAACTTTCCATCCTTTACTACAAATCAA |
| NOL4_18_bis_f | GGGAGGAGGAATTTTTAGGATTTTAT |
| NOL4_18_bis_r1 | AGTACCCATTATCCCATATCCTTCTCTAAAC |
| NOL4_18_bis_r2 | GATCGCCATTATCCCATATCCTTCTCTAAAC |
| PTPRN2_bis_f | AAGGAGAAGGGGGATATTAGTAGGTATT |
| PTPRN2_bis_r1 | AGTACCAATTCCTCCTTCAAAACAAACACT |
| PTPRN2_bis_r2 | GATCGCAATTCCTCCTTCAAAACAAACACT |
| SMARCA2_bis_f | TAGTTAGAGGGGAGAATGTTTAATGTG |
| SMARCA2_bis_r1 | AGTACATTACAAAAATAAATCACCACACCC |
| SMARCA2_bis_r2 | GATCGATTACAAAAATAAATCACCACACCC |

**Suppl. Figures**

Suppl. Fig. 1: Deep enzymology workflow allowing the methylation of DNA substrates in randomized $N_{10}CGN_{10}$ libraries, here shown for $N_5$ as example.

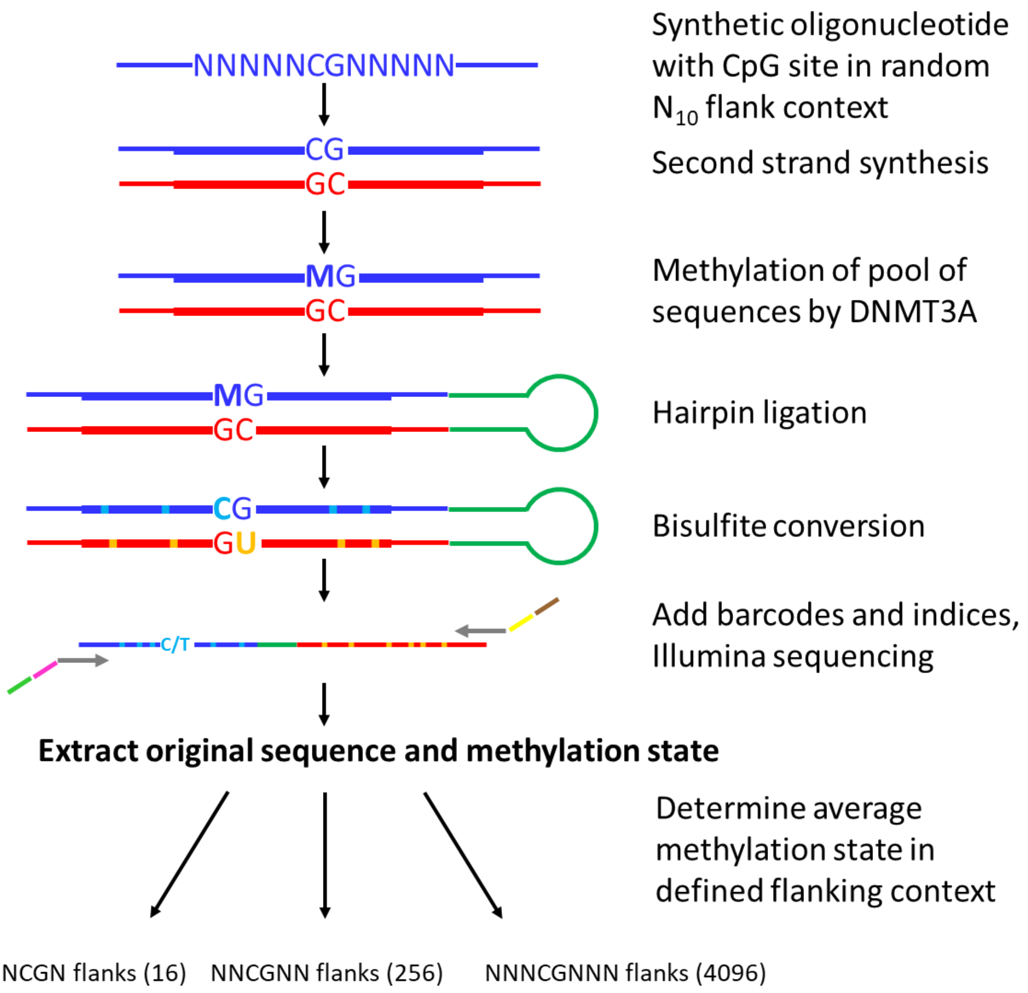

Suppl. Fig. 2: Correlation analysis of NNCGNN deep enzymology data obtained for DNMT3A, DNMT3B and R882H with hemimethylated (HM) and unmethylated (UM) substrates. For comparison, all data sets were reduced to equal read numbers of 70000. The figure shows Pearson correlation coefficients of the corresponding pairwise comparisons.

| | | DNMT3A | | DNMT3B | | R882H | |
|---|---|---|---|---|---|---|---|
| | | **HM** | **UM** | **HM** | **UM** | **HM** | **UM** |
| **DNMT3A** | **HM** | 1.00 | 0.92 | 0.34 | 0.29 | 0.68 | 0.65 |
| | **UM** | 0.92 | 1.00 | 0.34 | 0.31 | 0.65 | 0.61 |
| **DNMT3B** | **HM** | 0.34 | 0.34 | 1.00 | 0.89 | 0.74 | 0.60 |
| | **UM** | 0.29 | 0.31 | 0.89 | 1.00 | 0.69 | 0.67 |
| **R882H** | **HM** | 0.68 | 0.65 | 0.74 | 0.69 | 1.00 | 0.88 |
| | **UM** | 0.65 | 0.61 | 0.60 | 0.67 | 0.88 | 1.00 |

Suppl. Fig. 3: FCS profile of DNMT1 hypomorphic HCT116 cells with stable insertion of the DNMT3A (3A1) gene or its R882H mutant (3A1-R882H). DNMT3A expression was tested three days after induction by doxycycline. The vertical line indicates the threshold used for cell sorting in the following experiments.

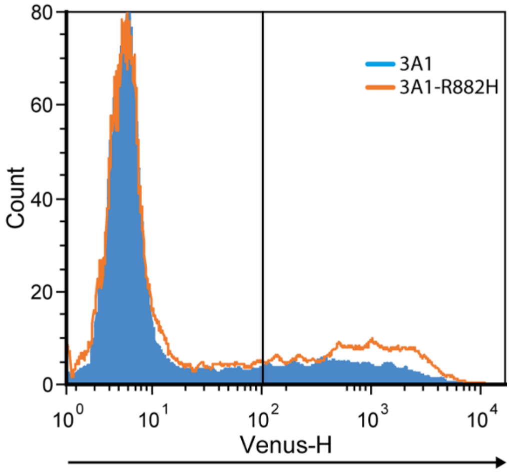

Supplemental Fig. 4: Definition of R882H/DNMT3A* preference profile as the average of the preferences of reverse complementary sites here illustrated for the example of N3 flanks and the AAA CG AAA and TTT CG TTT sites.

**Definition of R882H/DNMT3A* preference profile**

AAA CG AAA R882H/DNMT3A: 1.510
TTT CG TTT R882H/DNMT3A: 0.558
→ R882H/DNMT3A* preference for AAA CG AAA and TTT CG TTT: 1.034

Supplemental Fig. 5: Ranking of the flanks of CpG sites analyzed by Kim et al. (2013) to study major satellite methylation in murine ES cells in the R882H* preference (Kim et al., 2013). The figure displays the median, error bar denote the first and third median. Data were based on Southern Blot with MaeII (3 CpG sites) and bisulfite analysis (6 CpG sites). The majority of sites has a low preference for R882H as indicated by the high profile rank.

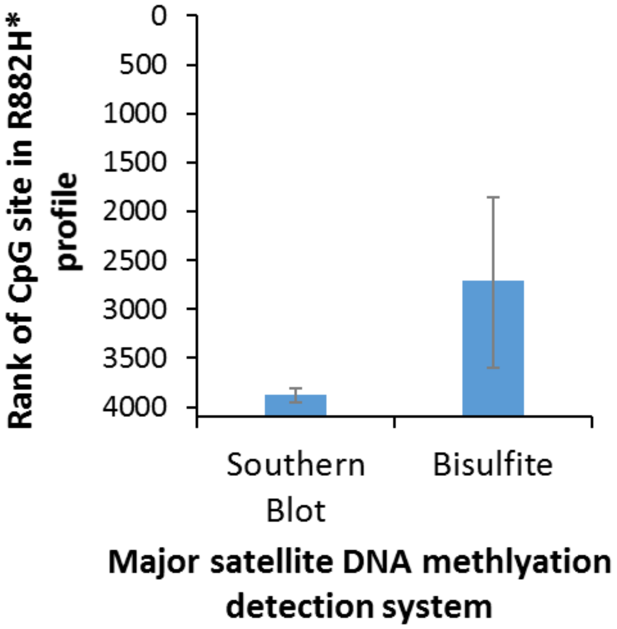

Suppl. Fig. 6: Binning of the 56580 sites that were specifically hypermethylated in R882H containing AML in the data set of Glass et al. (2017) into different tiers depending on their degree of hypermethylation (Glass et al., 2017).

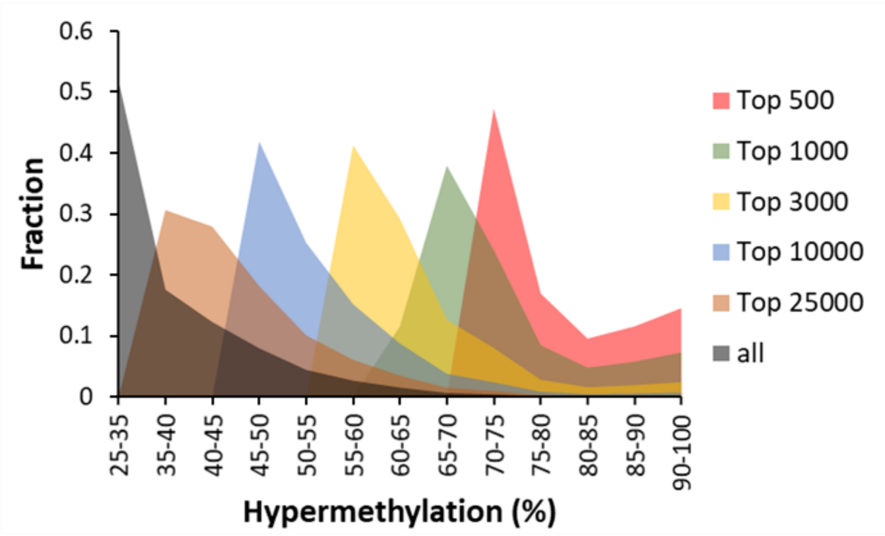
